# Precise topology of adjacent domain-general and sensory-biased regions in the human brain

**DOI:** 10.1101/2021.02.21.431622

**Authors:** Moataz Assem, Sneha Shashidhara, Matthew F. Glasser, John Duncan

## Abstract

Recent functional MRI studies identified sensory-biased regions across much of the association cortices and cerebellum. However, their anatomical relationship to multiple-demand (MD) regions, characterized as domain-general due to their co-activation during multiple cognitive demands, remains unclear. For a better anatomical delineation, we used multimodal MRI techniques of the Human Connectome Project to scan subjects performing visual and auditory versions of a working memory (WM) task. The contrast between hard and easy WM showed strong domain generality, with essentially identical patterns of cortical, subcortical and cerebellar MD activity for visual and auditory materials. In contrast, modality preferences were shown by contrasting easy WM with baseline; most MD regions showed visual preference while immediately adjacent to cortical MD regions, there were interleaved regions of both visual and auditory preference. The results may exemplify a general motif whereby domain-specific regions feed information into and out of an adjacent, integrative MD core.

## Introduction

The anatomical and functional organization of domain-general and domain-specific regions in the human brain remains unclear. On the one hand, thousands of functional MRI (fMRI) studies converge on a cortical, subcortical and cerebellar set of domain-general or multiple-demand (MD) regions that co-activate in association with many cognitively demanding tasks such as working memory, selective attention, and problem solving (Cole and Schneider 2007; Fedorenko et al. 2013; Hugdahl et al. 2015; Assem et al. 2020; Shashidhara et al. 2020). MD regions form a functionally integrated system, as revealed by the high correlations of their functional time-series during the presence or absence of a cognitive task (Power et al. 2011; Yeo et al. 2011; Blank et al. 2014; Ji et al. 2019; Assem et al. 2020; Cocuzza et al. 2020). MD’s fine-grained activation patterns flexibly change to reflect many types of task-relevant information (Woolgar et al. 2016), while in putative non-human primate (NHP) homologues, neurons respond to complex conjunctions of multiple task features (Rigotti et al. 2013; Stokes et al. 2013). These properties have suggested that MD regions play a central role in cognitive control, integrating the right information at the right time for the current cognitive operation (Miller and Cohen 2001; Cole and Schneider 2007; Duncan et al. 2020).

In a recent study, we utilized the Human Connectome Project’s (HCP) high quality multimodal MRI dataset and improved surface-based cortical alignment methods (Glasser, Smith, et al. 2016; Robinson et al. 2018) to better understand the anatomical and functional organization of the MD system (Assem et al. 2020). The conjunction of three cognitively demanding contrasts allowed us to delineate 9 specific MD patches per hemisphere, distributed in frontal, parietal and temporal association cortices. Subdividing each patch using the HCP’s recent multimodal cortical parcellation (HCP MMP1.0), we defined a core of 10 MMP regions that are most strongly activated and functionally interconnected, surrounded by a penumbra of 18 regions, which together we labelled as the extended MD system (**Figure 1a**). Though the MD system as a whole was co-activated by each contrast, improved anatomical specificity coupled with the statistical power of using hundreds of HCP subjects provided some of the strongest evidence in the literature indicating how each MD region has its own specific profile of *relative* activity across tasks (Assem et al. 2020). We suggested that broad MD co-activation reflects strong communication and integration between MD regions, while relative functional preferences reflect variations in local connectivity, and hence routes for many types of information to be fed into and out of the integrated MD system [(Duncan et al., 2020) see also (Power et al. 2013)].

**Figure 1.**
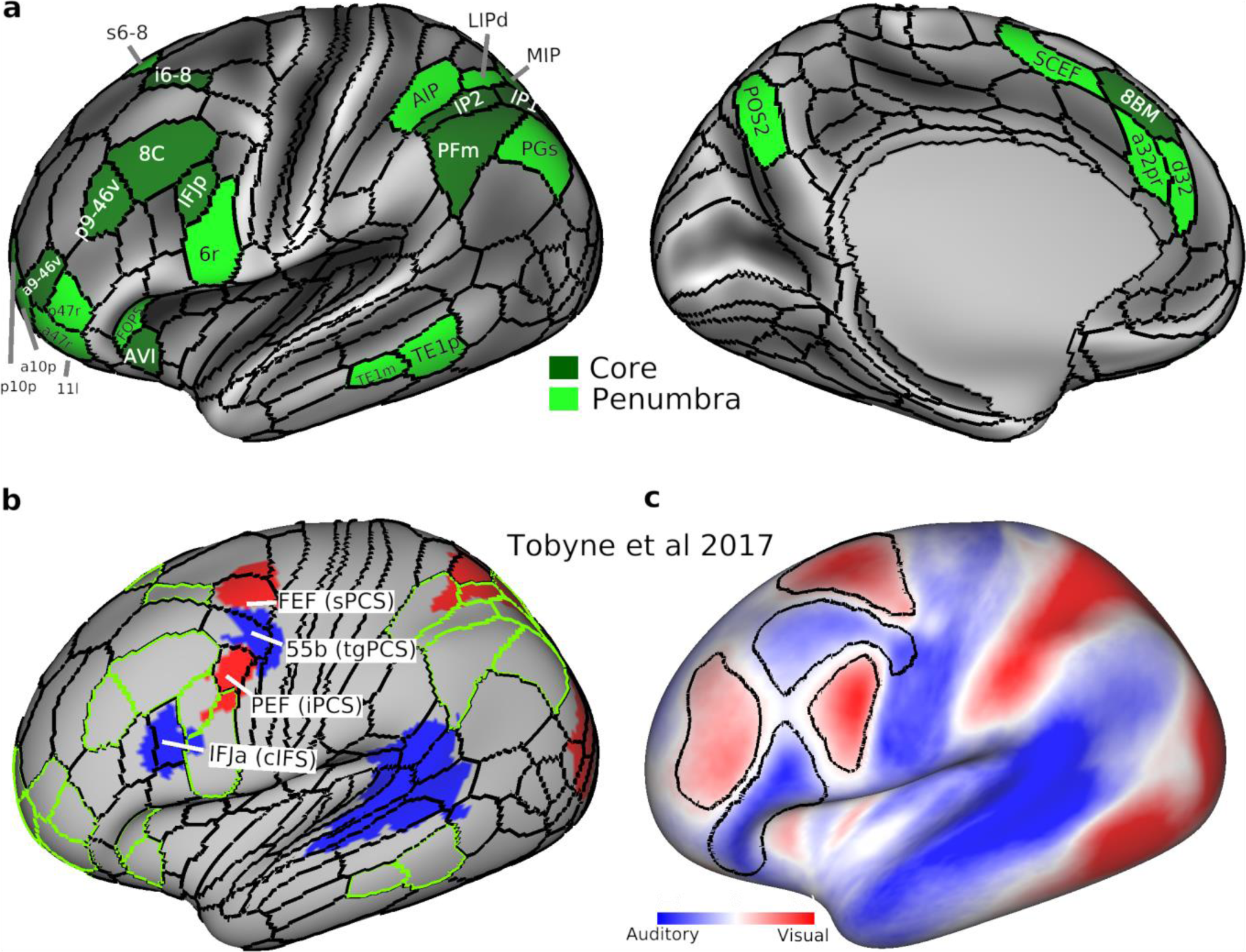
**(a)** The extended MD systemc. Core MD regions are colored in dark green and white labels. Penumbra MD regions are in light green with black labels. Note here we separated core region SCEF/8BM (as identified in Assem et al 2020) into SCEF as penumbra and 8BM as core for simplicity in analysis. **(b)** and **(c)** are adapted from Tobyne et al. (2017). **(b)** Sensory-biased regions [originally identified in (Michalka et al. 2015)] after their transformation to the HCP fs_LR surface. Red: visually biased. Blue: auditory biased. Overlapping HCP MMP1.0 regions are labelled and their original Michalka et al. (2015) labels are in brackets. Green contours correspond to extended MD borders in (a). **(c)** Sensory-biased lateral frontal regions based on their intrinsic rfMRI connectivity with posterior cortical areas. Black contours surrounding regions with warmer colours (red) are significantly more connected with visual parietal areas than auditory temporal regions. Black contours surrounding regions with colder colours (blue) are significantly more connected with auditory temporal regions than visual parietal regions. Data available at https://balsa.wustl.edu/6VPVv

Complementing this evidence for domain-generality, several anatomical, electrophysiological and recent fMRI studies robustly identified regions with sensory modality biases across the lateral frontal, parietal and temporal cortices (Romanski 2007; Michalka et al. 2015). More specifically, a recent series of fMRI studies contrasted visual and auditory attention or working memory tasks, along with careful examination of each individual’s cortical activations, to reveal four interdigitated lateral frontal regions showing relative sensory biases: two visual-biased regions along the superior and inferior precentral sulcus (sPCS and iPCS) interleaved with two auditory-biased regions, one in between the visual regions along the transverse gyrus (tgPCS) and another antero-ventral to the inferior visual region, often along the caudal portion of the inferior frontal sulcus (cIFS) (Michalka et al. 2015; Noyce et al. 2017; Lefco et al. 2020). More posteriorly, the same contrasts also identified visually biased parietal and auditory biased temporal regions. To estimate the overlap of these sensory-biased regions with the HCP MMP 1.0, Tobyne et al. (2017) applied a surface transformation approach. The four frontal interdigitated areas were found to overlap with FEF (visual), 55b (auditory), PEF (visual) and IFJa (auditory) all of which are, interestingly, just outside frontal MD regions (**Figure 1b**). To probe intrinsic frontal sensory biases beyond the previously mentioned four regions, Tobyne et al used the transformed parietal (visually biased) and temporal (auditory biased) regions as seeds to map out their functional connectivity with the lateral frontal cortex using HCP’s rfMRI data (with cortices aligned using MSMAll; see **Methods** section). The results revealed wider swathes of sensory-biased frontal regions (**Figure 1c**). Three visually preferring clusters were organized along caudal superior frontal sulcus, anterior middle frontal gyrus and inferior pre- central sulcus. These were interleaved with two auditory biased clusters, located along posterior middle frontal gyrus and inferior frontal sulcus (Tobyne et al. 2017).

One intriguing possibility is that sensory biased regions lie close to MD regions, in line with previous findings of side-by-side arrangement of domain-specific language and MD regions (Fedorenko et al. 2012; Fedorenko and Blank 2020). This could allow such regions to feed sensory-specific information into adjacent MD regions for integration into the current cognitive operation, perhaps exemplifying a general motif for the interaction between MD and adjacent, more domain-specific regions. Current data, however, leave unclear the precise relationship between domain-general and sensory-specific regions. While both sets of regions are currently available for direct comparison in the “HCP fs_LR cortical space”, the sensory-biased regions were localized using folding-based surface registration, a process prone to anatomical mis- estimations due to variability of cortical folds across individuals (Coalson et al. 2018). Further, the large swathes of frontal sensory biases revealed using rfMRI (Tobyne et al. 2017) (**Figure 1c**) were identified using large parietal and temporal seed regions that likely mix signals from several functionally distinct regions (Glasser, Coalson, et al. 2016). The relationship between sensory biases and MD regions outside the lateral frontal lobe is also uncertain. For example, one study found that localized anterior cingulate and insular regions showed no sensory biases when contrasting a visual with an auditory working memory task (Noyce et al. 2017). This is in contrast to another study which identified a caudal (auditory) to rostral (visual) gradient throughout lateral, medial frontal and medial parietal cortices (Mayer et al. 2016). Finally, the functional properties of recently identified subcortical and cerebellar MD regions (Assem et al. 2020) remain uncharted territories. Some fMRI studies have failed to find subcortical modality preferences (Bushara et al. 1999; Mayer et al. 2016), while in the cerebellum, evidence for visual and auditory responses was reported in Cruses I and II (Petacchi et al. 2005; Kirschen et al. 2010; Brissenden et al. 2018; Ren et al. 2021), where MD cerebellar regions have also been localized (Assem et al. 2020).

To resolve these questions with a high anatomical accuracy, we collected a new dataset using HCP’s multimodal MRI acquisition and analysis approaches, which utilize surface based approaches and multimodal MRI features for accurate alignment of cortical regions across individuals (Glasser, Smith, et al. 2016; Robinson et al. 2018). As recently demonstrated, traditional brain imaging approaches will miss out on robust evidence of sensory-biased regions (Noyce et al. 2017; Lefco et al. 2020) due to their reliance on suboptimal methods (e.g. unconstrained volumetric smoothing and 3D volumetric alignment) which are inherently inferior to surface-based methods and fail to accurately align many individual differences in areal topographies (Coalson et al. 2018).

To probe MD activity for this study, we chose a working memory (WM) paradigm as an example of a cognitive demand that is well recognized to activate MD regions (Fedorenko et al. 2013; Assem et al. 2020). The same subjects performed visual (day 1) and auditory (day 2) versions of the n-back WM task, with each modality having two difficulty levels (easy and hard).

This is a critical manipulation because MD regions are characterized by their strong activations to task difficulty manipulations (Fedorenko et al. 2013; Assem et al. 2020), but none of the previous studies probed the interaction between MD difficulty and sensory preferences (Michalka et al. 2015; Noyce et al. 2017).

## Materials and Methods

### Subjects

Thirty-seven human subjects participated in this study (age=25.9±4.7, 23 females, all right-handed). Originally fifty subjects were scanned over two sessions; thirteen subjects were excluded due to incomplete data (n=5), excessive head movement during scanning (n=4), or technical problems during scanning (n=2) or analysis (n=2). All subjects had normal or corrected vision (using MRI compatible glasses). Informed consent was obtained from each subject and the study was approved by the Cambridge Psychology Research Ethics Committee.

### Task Paradigms

Each subject performed five tasks in the scanner over two sessions. The current study used data from two tasks: visual n-back (session 1), auditory n-back (session 2) (**Figure 2**). Each n-back task was performed for four runs, and each run consisted of four 1-back (easy) and four 3- back (hard) blocks. Each task block (30 s) started with a cue (4 s) followed by 12 trials (24 s, 2 s each) and ended with a blank screen (2 s) as an inter-block interval. Task blocks were paired (easy followed by hard, or hard followed by easy) and the order was counterbalanced across runs and subjects. A fixation block (16 s) followed every two paired task blocks. In the visual session, each run consisted of 36 blocks: 8 visual n-back blocks, 12 fixation blocks, and 8 blocks for each of two other visual tasks. In the auditory session, each run consisted of 8 auditory n-back and 4 fixation blocks. In the auditory session, n-back runs were alternated with runs of another visual task not analysed here.

**Figure 2.**
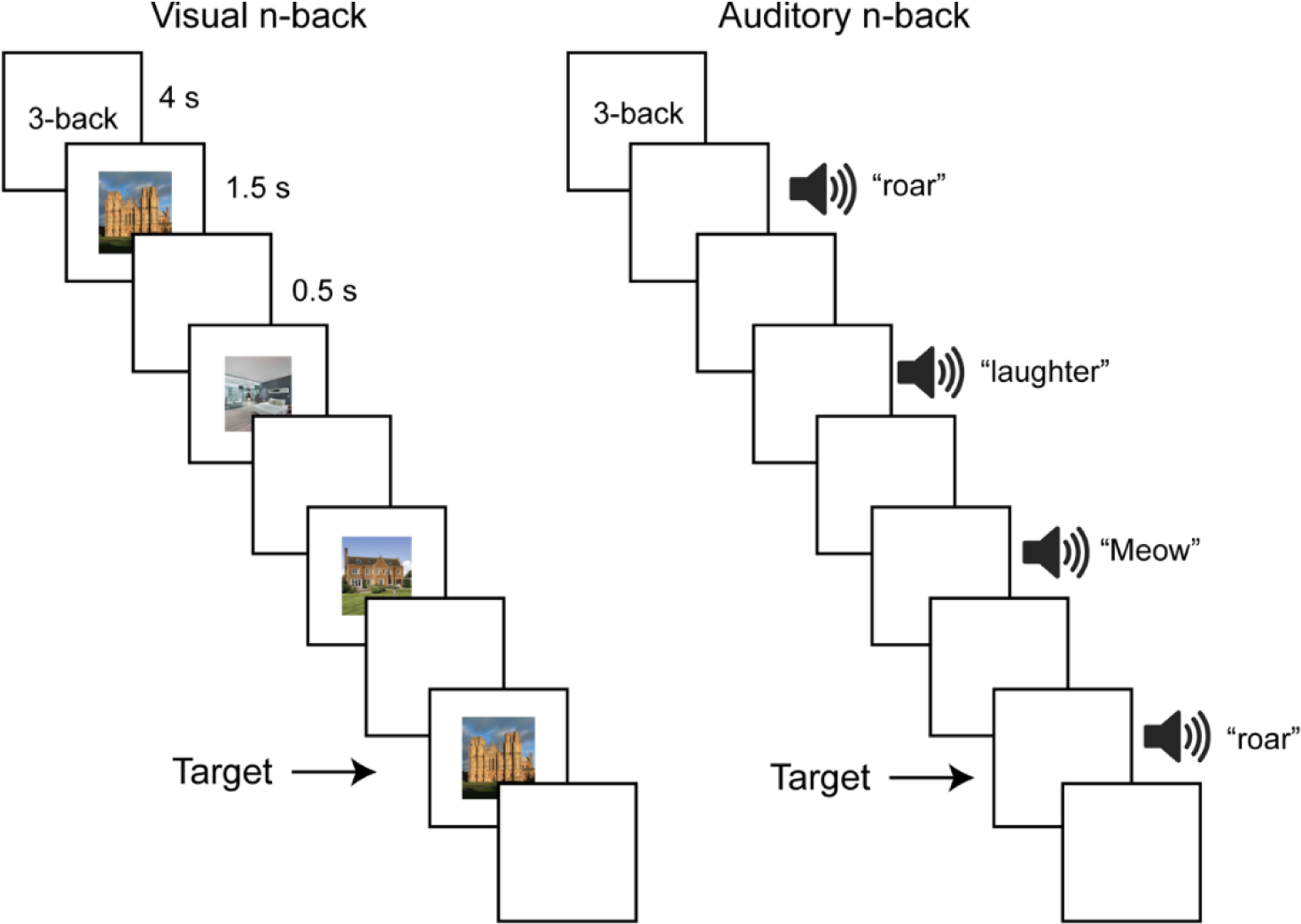
N-back task paradigm. Illustration of a stimulus sequence from the hard (3-back) version. Subjects performed a visual and an auditory version of the n-back task in separate sessions. Each task had easy (1-back, not shown) and hard (3-back) blocks. Each block (30 s) started with a cue (4 s) followed by 12 trials (2 s each) and ended with a fixation screen (2 s). Subjects pressed right for target stimuli, and left for all non-target stimuli. For the visual n-back, stimuli consisted of pictures of houses (illustrated) or faces (not shown). Auditory n-back stimuli consisted of animate (illustrated) or inanimate (not shown) sounds. See **Materials and Methods** for further details.

Each trial lasted for 2 s. The visual stimulus was presented for 1500 ms, followed by 500 ms of a blank screen. Auditory stimuli had a duration of 1250 ms (except for two sounds which were 1360 and 1520 ms long), followed by 480-750 ms of a blank screen. Responses were accepted at any moment throughout the trial. For the 3-back condition, subjects were instructed to press right for the target stimulus (i.e. current stimulus was the same as the one 3 steps back), and left for all non-target presentations. Similarly, for the 1-back condition, subjects were instructed to press right for the target stimulus (i.e. current stimulus was an exact repetition of the immediate previous stimulus) and press left for all non-target stimuli. In each block there were 1-2 targets. For the visual n-back, stimuli consisted of pictures of faces and houses. Face stimuli were selected from the Developmental Emotional Faces Stimulus Set (Meuwissen et al. 2017). Faces were either males or females, children or adults, making a happy or sad face. House stimuli were pictures of houses or churches, old or new, from inside or outside. There were 32 faces and 32 houses, each made up of 4 examples for each of the 2 x 2 x 2 possible feature combinations. These categories were necessary for other visual tasks during the session and have no bearing here. Auditory n-back stimuli consisted of animate (e.g. a human’s cough, a lion’s roar) and inanimate (e.g. a musical instrument, a bell ringing) sounds. There were 9 animate and 9 inanimate sounds. Faces and houses were presented in separate blocks, as were animate and inanimate sounds. Subjects were encouraged to use their right hand and respond to targets using a middle finger press and to non-targets using an index finger press but this was not enforced and several subjects found it more comfortable to use both hands for responses (index fingers or thumbs). Similarly, during the auditory task, subjects kept their eyes open or closed according to their preference.

### Image Acquisition

Images were acquired using a 3T Siemens Prisma scanner with a 32-channel RF receive head coil. MRI CCF acquisition protocols for HCP Young Adult cohort were used (package date 2016.07.14; https://protocols.humanconnectome.org/CCF/). These protocols are substantially similar to those described in previous studies (Glasser et al. 2013; Smith et al. 2013; Uǧurbil et al. 2013) but do differ in some respects. All subjects underwent the following scans over two sessions: structural (at least one 3D T1w MPRAGE and one 3D T2w SPACE scan at 0.8-mm isotropic resolution), rest fMRI (2 runs × 15 min), and task fMRI (5 tasks, 4 runs each, approx. 100 min total). Whole-brain rest and task fMRI data were acquired using identical multi-band (factor 8) EPI sequence parameters of 2-mm isotropic resolution (TR = 800 ms, TE=37 ms). Both rest and task EPI runs were acquired in pairs of reversed phase-encoding directions (AP/PA). Spin echo phase reversed images (AP/PA) were acquired during the structural and functional (after every 2 functional runs) scanning sessions to (1) correct fMRI images for phase encoding direction EPI distortion, (2) correct T1w and T2w images for readout distortion, (3) enable accurate cross-modal registrations of the T2w and fMRI images to the T1w image in each subject, (4) compute a more accurate fMRI bias field correction and (5) segment regions of gradient echo signal loss.

### Data preprocessing

Data preprocessing was also substantially similar to the HCP’s minimal preprocessing pipelines detailed previously (Glasser et al. 2013). A brief overview and differences are noted here. HCP pipelines versions 3.27.0 were used (scripts available at: https://github.com/Washington-University/HCPpipelines). For each subject, structural images (T1w and T2w) were used for extraction of cortical surfaces and segmentation of subcortical structures. Functional images (rest and task) were mapped from volume to surface space and combined with subcortical data in volume to form the standard CIFTI grayordinates space. Data were smoothed by a 2mm FWHM kernel in the grayordinate space that avoids mixing data across gyral banks for surface data and avoids mixing areal borders for subcortical data.

From this point onwards HCP pipelines version 4.0.0 were used (also available through the link above; specific parameters different from the default values are noted below). Rest and task fMRI data were additionally identically cleaned for spatially specific noise, largely from head motion, using spatial ICA+FIX (Salimi-Khorshidi et al. 2014). ICA+FIX was applied separately to each of the following concatenated runs: resting-state runs (2x15 mins), visual runs from session one (4x15 mins), and auditory runs (4x5 mins). Note the visual runs were longer because they included other tasks that are irrelevant to this analysis. An improved FIX classifier was used (provided by M.F.G.) for more accurate classification of noise components in task fMRI datasets. After manual checking of ICA+FIX outputs for 10 subjects, a threshold of 50 was determined for “good” vs “bad” signal classification and applied for the remaining subjects. In contrast to the Assem et al. (2020) study, global structured noise, largely from respiration, was not removed using temporal ICA as public scripts are not yet available.

For accurate cross-subject registration of cortical surfaces, the multimodal surface matching algorithm MSM was used. First “sulc” cortical folding maps are gently registered in the MSMSulc registration, optimizing for functional alignment without overfitting folds. Second, a combination of myelin, resting-state network, and rest fMRI visuotopic maps (Robinson et al. 2014, 2018) was used to fully functionally align the data. For this purpose, we used 30 mins of resting state data, acquired in the second session prior to the auditory task.

### Task fMRI analysis

Task fMRI analysis scripts in HCP pipelines version 4.0.0 were used. Default steps are detailed in Barch et al. (2013). Briefly, autocorrelation was estimated using FSL’s FILM on the surface (default parameters in the HCP’s task fMRI analysis scripts were used). Activation estimates were computed for the preprocessed functional time series from each run using a general linear model (GLM) implemented in FSL’s FILM (Woolrich et al. 2001).

For each of the n-back tasks, 4 regressors were used (2 stimulus category x 2 task difficulty). Each predictor had a unitary height and covered the period from the onset of the cue to the offset of the final trial (28 sec). All regressors were then convolved with a canonical hemodynamic response function and its temporal derivative. 12 additional motion regressors were added to the model (3 translation, 3 rotation and their derivatives). The time series and the GLM design were temporally filtered with a Gaussian-weighted linear highpass filter with a cutoff of 200 seconds. Finally, the time series was prewhitened within FILM to correct for autocorrelations in the fMRI data. Surface-based autocorrelation estimate smoothing was incorporated into FSL’s FILM at a sigma of 5mm. Fixed-effects analyses were conducted using FSL’s FEAT to estimate the average effects across runs within each subject.

For further analysis of effect sizes, beta ‘cope’ maps were generated using custom built MATLAB scripts after moving the data from the CIFTI file format to the MATLAB workspace. Beta maps were then converted to percent signal change as follows: 100*(beta/10000). The value 10000 corresponds to the grand mean scaling of the overall timeseries of each run during preprocessing. Unless mentioned otherwise, parametric statistical tests were used.

For parcellating the cerebral cortex, the group-average HCP multi-modal parcellation (MMP1.0) was used (Glasser, Coalson, et al. 2016), as the individual-specific areal classifier is not publicly available. Still, due to the superior cortical alignment approach of MSMAll, the fraction of the individually defined regions that are captured by group-defined borders reaches 60–70% (Coalson et al. 2018) and we have previously demonstrated that comparing areal classifier and group-defined borders showed similar results (Assem et al. 2020). Values of vertices sharing the same areal label were averaged together to obtain a single value for each area.

The gradient map was created using wb_command –cifti-gradient function from the connectome workbench with a pre-smoothing sigma of 1 mm and using subject specific vertex areas and the midthickness cortical surface.

For subcortical and cerebellar analysis, an MD mask covering regions of the caudate, thalamus and cerebellum was used. In Assem et al (2020) two versions of the subcortical/cerebellar MD masks were defined: One based on a conjunction of task activations and one based on rfMRI connectivity with cortical MD core. In this study, the mask based on rfMRI was utilized because (1) it includes putative thalamic MD regions that are not included in the task-based mask (2) task and rest fMRI masks show substantial overlap in the remaining caudate and cerebellar regions. The volumetric cerebellar results were projected on a flat cerebellar surface using SUIT software (Diedrichsen and Zotow 2015).

## Results

Thirty-seven subjects were scanned while performing visual and auditory n-back tasks, with each modality in a separate session. Each n-back task had an easy (1-back) and a hard (3-back) version (**Figure 2**; see **Methods**). Structural and fMRI data were preprocessed using the HCP pipelines. Cortical surfaces were functionally aligned across individuals using multimodal MRI features (MSMAll), the HCP’s multi-modal parcellation version 1.0 was used for defining cortical areas, and subcortical and cerebellar regions were extracted separately for each individual (see **Methods**).

### Behaviour

As expected, across four runs for each task, performance on the easy condition was better than the hard condition for both visual (see **Table 1;** accuracy t(36):8.4, p<10^-9^; reaction time (RT) t(36):14.1 p<10^-15^) and auditory (accuracy t(36):8.9, p<10^-9^; RT t(36):1.7, p=0.1) tasks. Subjects were more accurate on the visual than the auditory task during both the easy (t(36): 3.2, p<0.01) and hard conditions (t(36): 5.1, p<10^-5^). Any differences in RTs between visual and auditory conditions would be uninterpretable as the auditory stimulus took a longer time to be presented (see **Methods**). Lastly, as expected, accuracy on non-target (NT) trials was better than target (T) trials for both visual (easy t(36):4.2, p<0.001; hard T<NT: t(36):8.8, p<10^-9^) and auditory (easy T<NT t(36):-6.4, p<10^-6^; hard T<NT: t(36):-12.3, p<10^-13^) tasks.

**Table 1.** Behavioural performance.

|  | **Visual** |  | **Auditory** |  |
| --- | --- | --- | --- | --- |
|  | Easy | Hard | Easy | Hard |
| % Correct<br>(all trials) | 95.5±5.1 | 86.7±5.7 | 92.1±6.6 | 78.4±8.8 |
| RT (s)<br>(all trials) | 0.61±0.08 | 0.79±0.1 | 1.31±0.1 | 1.33±0.1 |
| % Correct<br>(Target trials) | 92.5±9.2 | 78.8±10.6 | 86.2±12 | 67.2±13.7 |
| % Correct<br>(Non-Target trials) | 98.4±2 | 94.5±3.2 | 97.9±2.4 | 89.6±5.5 |

### MD cortex visual vs auditory activations during a task difficulty manipulation

We first sought to investigate cortical sensory modality biases for the hard>easy contrast. As demonstrated in previous studies, this difficulty manipulation is a strong driver of MD co- activations (Fedorenko et al. 2013). **Figure 3a** shows average MD activations for three cognitively demanding contrasts (WM, reasoning and math) from our previous study (Assem et al. 2020) with the green contours highlighting the extended MD regions.

**Figure 3.**
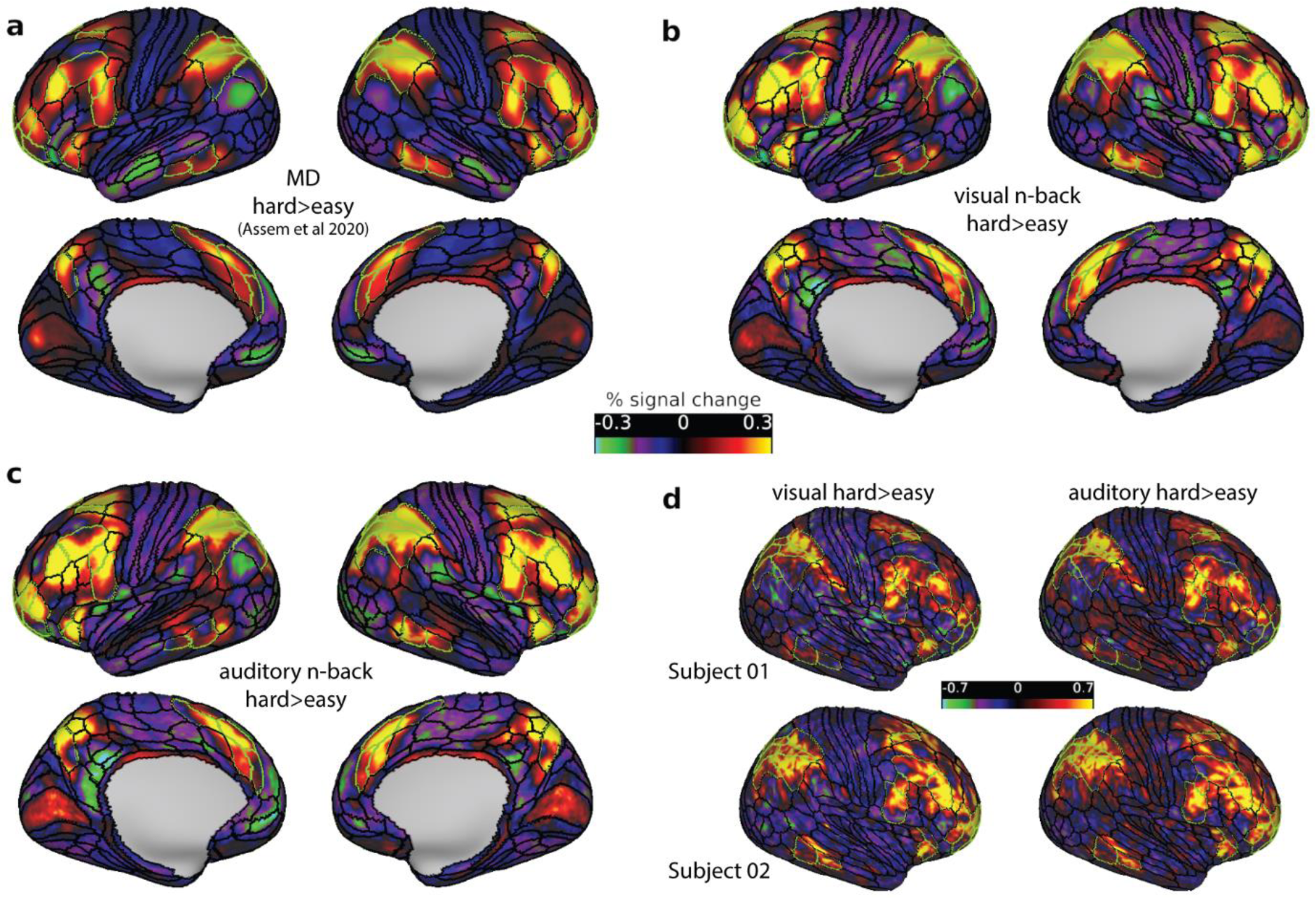
Hard>easy n-back activation maps. (a) MD group (n=449) average activation of three cognitively demanding contrasts from Assem et al. (2020). (b) Visual and (c) auditory group (n=37) average n-back hard>easy activations. (d) Two exemplar single subject activation maps for the hard>easy visual (left) and auditory (right) n-back task. All activation values are percent signal change. Data available at https://balsa.wustl.edu/1BkBG and all 37 subject activation files can be downloaded from https://balsa.wustl.edu/5XxX1

For an initial comparison, we examined the group level activations for the visual and auditory modalities separately (**Figure 3b and 3c).** We averaged activation maps for each modality across the four runs due to their high correlations (minimum whole brain Pearson’s *r* between any two runs=0.90). MSMAll registration significantly improves the alignment of areas across subjects with peak probability overlaps reaching >90% for most areas (Coalson et al. 2018) thus allowing us to identify activations overlapping with MD areas. As expected, the hard>easy activations showed substantial similarity with the average MD activations in **Figure 3a** (correlation between MD and hard>easy visual *r*=0.86, MD and hard>easy auditory *r*=0.84) and peak activations overlapped with extended MD borders (green contours). Unexpected, though, is the striking similarity between the visual and auditory hard>easy contrasts (correlation of all cortical vertices activations between both maps *r*=0.96; and between MD vertices only *r*=0.98). This similarity was not an artefact of averaging activations across subjects as it was also evident in individual subject activation maps [average (for all cortical vertices) mean *r*=0.71, range 0.43-0.82] (**Figure 3d**).

To quantify hard>easy activations across the 28 cortical areas classified as parts of the extended MD regions for each hemisphere (**Figure 1a**), for each area we averaged the activation estimates for all vertices and performed a one-sample t-test across subjects against a mean of zero. For this analysis, we have also included the 4 MMP1.0 interdigitated regions as approximations of the Michalka et al regions: visual-biased: FEF, PEF, auditory-biased: 55b and IFJa (**Figure 1b**). As expected, extended MD regions showed significant activation (**Figure 4;** p<0.05, Bonferroni corrected for n=64 regions) for both modalities except for one region in the right hemisphere (SCEF) and four regions in the left hemisphere: a47r, PGS, TE1p, and TE1m. One left region (d32) was only active for the visual contrast. These results show stronger MD activations in the right hemisphere (t(36)=2.15, p=0.038), replicating the findings of the visual n-back task in our previous study (Assem et al. 2020). As for the Michalka regions, the visually biased FEF and PEF were significantly activated for both the visual and auditory tasks in both hemispheres (**Figure 4;** p<0.05, Bonferroni corrected for n=64 regions). However, auditory biased regions showed mixed results: IFJa showed significant activations in both hemispheres for the auditory contrast but only left side significant activations for the visual contrast. Area 55b showed significant activations for the auditory contrast in the left hemisphere only (and the right hemisphere at a lower Bonferroni threshold n=4 regions).

**Figure 4.**
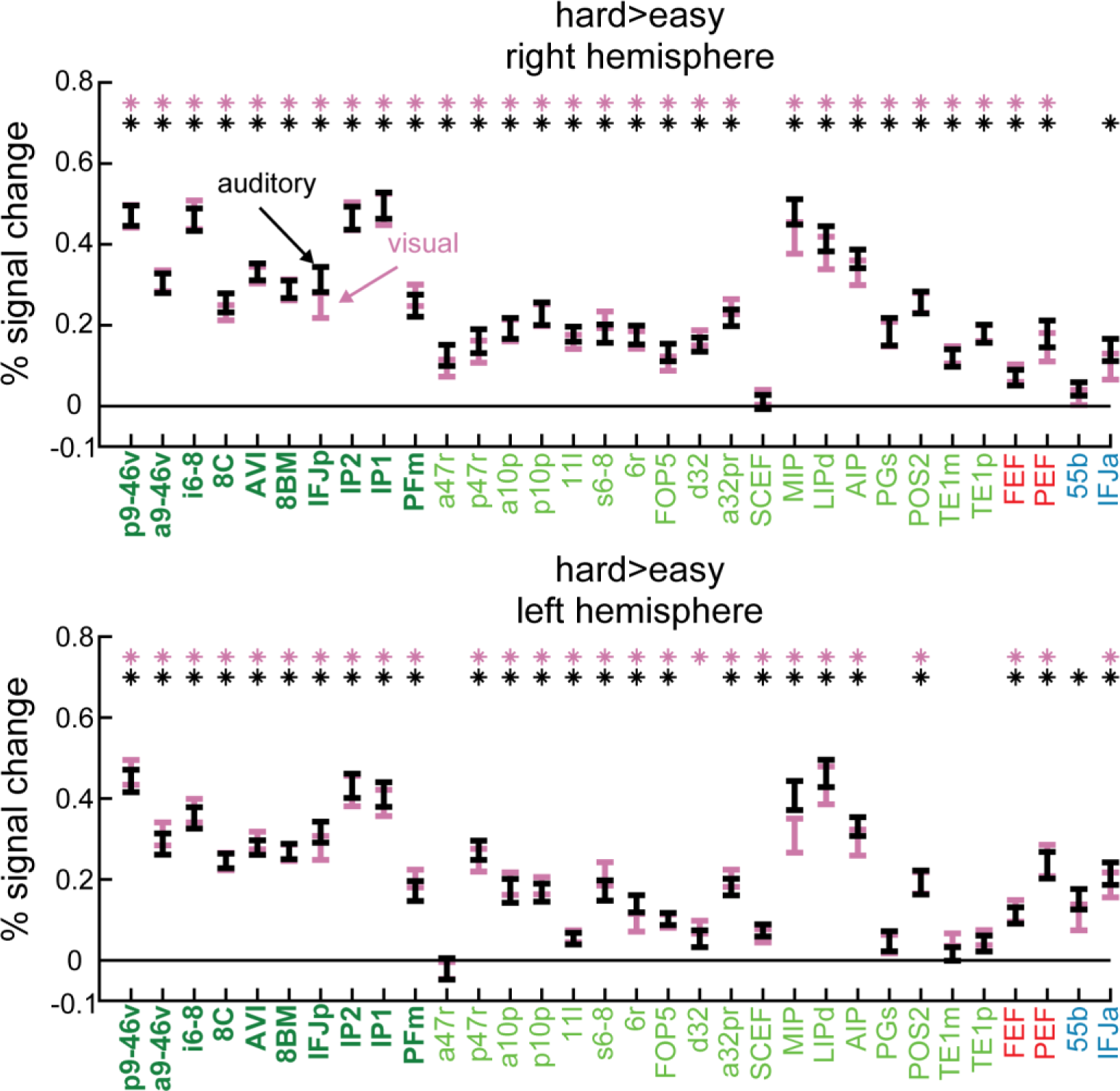
Hard>easy contrast activations (% signal change) in right (top) and left (bottom) hemispheres, separately for auditory (black) and visual (pink) tasks. Error bars represent standard error of the mean (SEM). Extended MD regions labels are coloured in green (core MD in dark green and bold, penumbra MD in light green). The 4 MMP1.0 regions partially overlapping with Michalka et al (2015) sensory-biased regions are coloured in red (visual-biased) and blue (auditory-biased). Asterisks denote p<0.05 Bonferroni corrected for n=64 regions.

For completeness, we also performed a conjunction analysis for all cortical regions significantly activated (p<0.05, Bonferroni corrected for 360 regions) in both the visual and auditory hard>easy contrast. The results in **Supplementary figure 1** confirm that a total of 27 non-MD and non-Michalka et al regions (across both hemispheres) are activated in both task contrasts. These additional regions replicate our previous findings (Assem et al. 2020) of more domain-specific activations accompanying domain-general MD activity, in this case possibly reflecting specific requirements of the n-back task.

Overall, these results confirm that the hard>easy manipulation in the n-back task engaged extended MD regions and that these regions are activated by both visual and auditory modalities. Previously identified sensory biased regions were also engaged, though not always significantly for both modalities.

Next, to investigate MD sensory preferences, for each subject we subtracted the auditory hard>easy map from the visual hard>easy map and extracted a single value for each region by averaging across its vertices. Then for each region we performed a one-sample t-test across subjects against zero. We failed to find any significant difference between modalities across both extended MD regions and the MMP1.0 approximations of Michalka et al (2015) sensory biased regions (i.e. FEF, 55b, PEF and IFJa), in either hemisphere (**Figure 5a;** p<0.05, Bonferroni corrected for n=64 regions). There was also no significance for the Michalka regions with a less conservative Bonferroni correction (n=4 regions).

**Figure 5.**
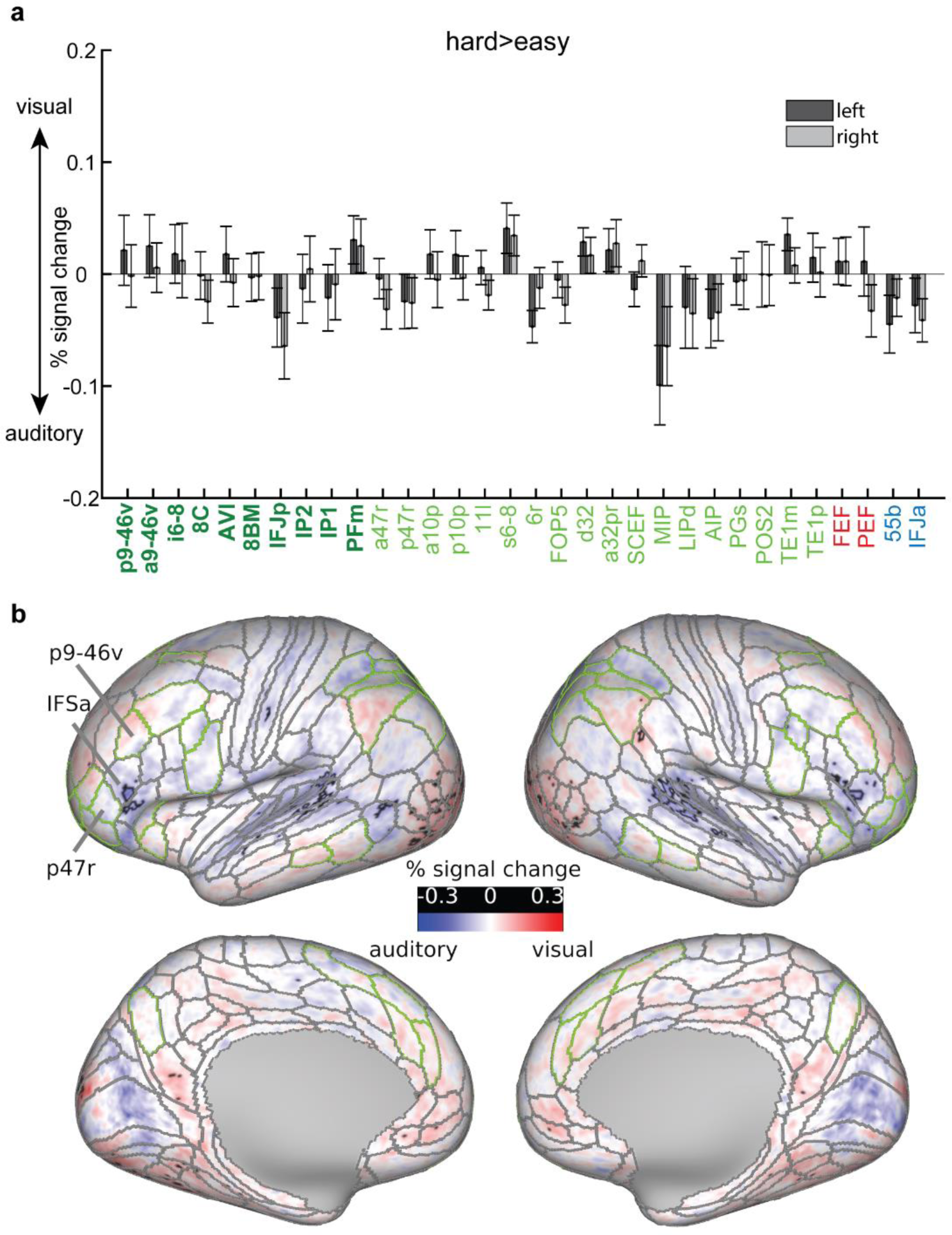
Visual vs auditory task preferences for the hard>easy contrast. **(a)** Bar heights represent average activation differences (%signal change, visual hard>easy minus auditory hard>easy) for each region across subjects. Error bars represent SEM. Light coloured bars represent regions of the right hemisphere. Areal names are coloured in dark bold green (core MD), light green (penumbra MD), red (Michalka-visual), blue (Michalka-auditory) **(b)** Cortical activations (% signal change) for the same contrast. Black contours surround significant vertices (FDR corrected p<0.05), grey contours correspond to the HCP MMP 1.0 areal borders and green contours correspond to extended MD areal borders. Data available at https://balsa.wustl.edu/npmpr

To uncover potential finer grained regions with stronger visual or auditory activations, we repeated the one-sample t-test on each cortical vertex (FDR corrected p<0.05). This analysis again failed to identify any contiguous sets of significant vertices within extended MD regions (**Figure 5b**). We identified a small bilateral set of vertices overlapping with IFSa that showed stronger activations during the auditory task (**Figure 5b**). This region is more anterior than previously reported frontal auditory biased regions (Michalka et al. 2015; Tobyne et al. 2017) and lies in between two MD regions p9-46v (caudal) and p47r (rostral). Unsurprisingly, we also identified small groups of significant vertices that overlapped with early auditory regions and visual extrastriate regions (**Figure 5b**).

To explore individual differences in cortical activations, we identified a group of 6 subjects with a mean auditory task accuracy better than the visual task during the hard condition, contrasting with the 31 subjects with the reverse pattern. However, none of the extended MD or Michalka vertices or regions for either group showed a significant auditory vs visual task bias (p<0.05 FDR corrected for vertex-wise analysis, Bonferroni corrected n = 64 for region-wise analysis, or just n=4 for the Michalka regions).

Taken together, these results highlight that while MD cortex increased its activity during a demanding WM manipulation, it showed no statistically significant preference for either the visual or auditory hard>easy contrast. This contrast also failed to identify the previously reported interdigitated pattern of sensory biases in the frontal cortex (**Figure 1 b and 1c**). However, we did identify a novel anterior ventral frontal region (IFSa) with stronger activation for the auditory hard>easy contrast.

### Visual vs auditory activations during low cognitive demands

Why did the hard>easy contrast fail to replicate sensory biases, robustly identified in previous studies, across much of the frontal and parietal cortices (Michalka et al. 2015; Noyce et al. 2017; Tobyne et al. 2017)? In an attempt to reproduce these sensory-biased regions, we sought to investigate visual and auditory activations for the easy>fix contrast. One possibility could be that hard>easy activations, similar for the two modalities, add to a background of sensory bias already existing in the easy tasks.

Hence, we repeated the same analysis in the previous section using the easy>fix contrast. For each subject we subtracted the visual easy>fix map from the auditory easy>fix map (**Figure 6a)** then we performed a one sample t-test across subjects for each vertex (FDR corrected p<0.05; **Supplementary figure 2**). On the lateral frontal surface, the contrast activations now highlight an interdigitated pattern of visual vs auditory task biases similar to previous reports though with a crisper anatomical delineation (**Figure 6a**). In line with Tobyne et al.’s estimation, FEF and PEF showed stronger activations during the visual than auditory task. In between FEF and PEF, a small region in the posterior portion of 55b showed stronger activations during the auditory contrast (**Figure 6a).** Stronger visual task activations in PEF extended anteriorly towards IFJp and IFJa. Within IFJa, we found that its dorsal segment had stronger visual task activations, while its ventral segment had stronger auditory task activations. This division was more prominent in the left hemisphere (**Figure 6a**). Further anteriorly, we identified two more interdigitating regions. IFSp had stronger visual task activations, in line with previous indications of a new anterior visually-biased region (Lefco et al. 2020). More anteriorly, IFSa had stronger auditory task activations, matching our hard>easy findings in the previous section. Even more anteriorly near the frontal pole, we identify a patch with stronger visual task activations mostly overlapping with p47r (penumbra MD), just ventral to core MD region a9-46v.

**Figure 6.**
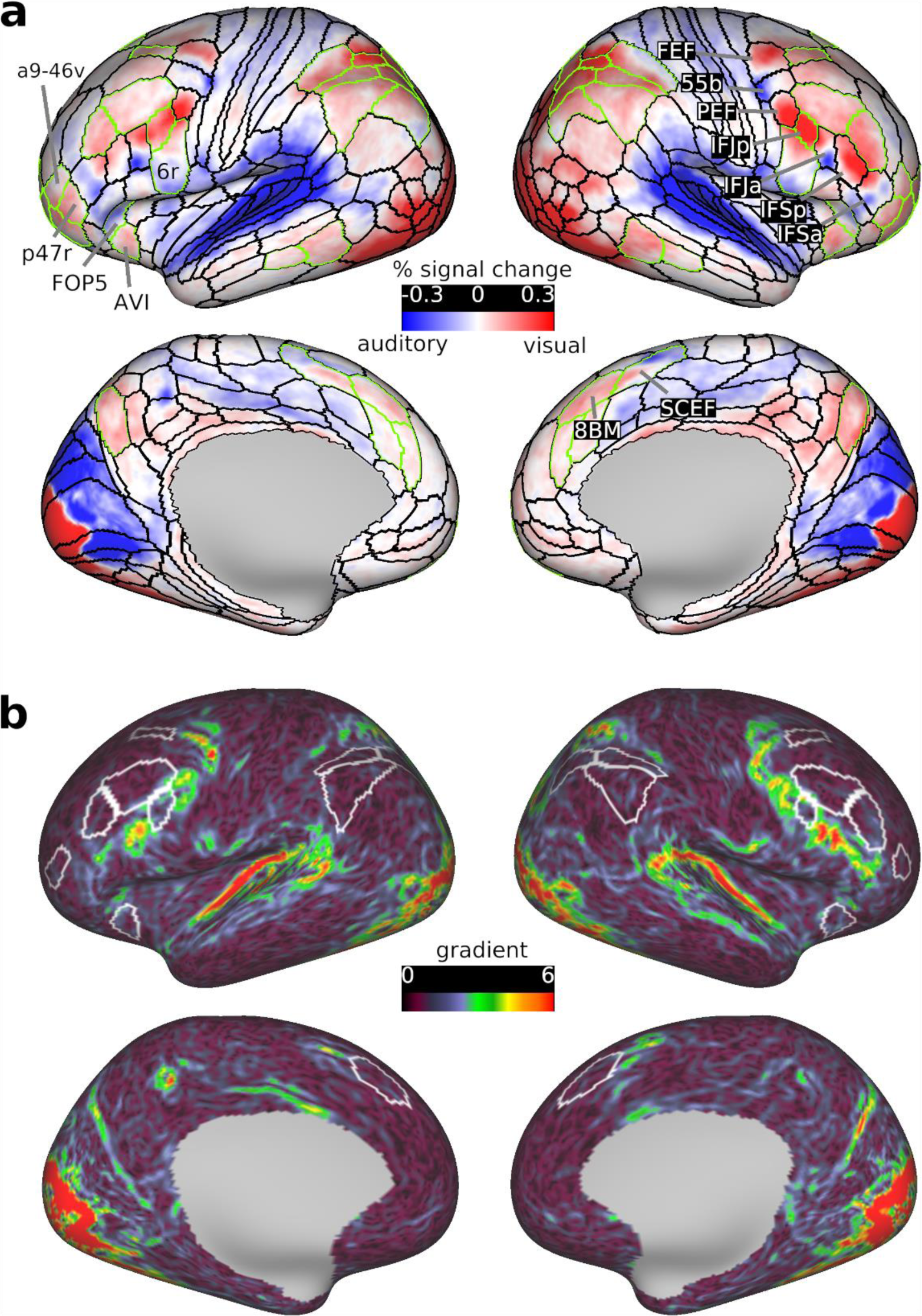
Visual vs auditory task preferences for the easy>fix contrast. **(a)** Cortical activations (%signal change) for the visual easy>fix minus auditory easy>fix contrast. **(b)** Gradient map (i.e. 1^st^ spatial derivative) for this contrast. Warmer colors highlight cortical regions with sudden shifts in task modality preferences. Note how the strongest gradients lie just outside core MD regions (white borders). Data available at https://balsa.wustl.edu/g7V7B including all 37 subject activation files for the contrast in **(a)**

Importantly, these interdigitations would not be visible at the group level using volumetric or even conventional surface-based inter-subject cortical alignment approaches (Noyce et al. 2017). This necessitated previous studies to use individual subject localizer approaches to uncover the interdigitations (Fedorenko et al. 2012; Nieto-Castañón and Fedorenko 2012; Michalka et al. 2015; Noyce et al. 2017). Here, however, the superior alignment approach of MSMAll uncovered this fine grained functional organization, just as predicted by (Coalson et al. 2018) and despite the frontal premotor regions having one of the highest inter-subject variability rates in areal topographies (Glasser, Coalson, et al. 2016).

On the lateral frontal surface, the peak activations of interdigitated visual and auditory preference lie along an arc posterior and ventral to frontal MD regions, crossing MD at IFJp (**Figure 6a**). To probe this pattern further, we computed a gradient map for the easy>fix visual minus easy>fix auditory activation map. A gradient map is akin to the first spatial derivative of the activation map (see **Methods**). Vertices with higher gradient values reflect a rapid shift in activation magnitudes, relative to all its neighbouring vertices [see supplementary methods in (Glasser, Coalson, et al. 2016)]. **Figure 6b** indeed shows that prominent gradients, which reflect rapid shifts in sensory preference, lie just adjacent to frontal core MD regions i6-8, 8C and p9- 46v. A similar gradient was observed outside parietal MD regions IP1 and IP2 and medial frontal region 8BM. Together, the gradient and activation maps show that strong visual vs auditory task biases occur just adjacent to frontal and parietal MD regions.

To more thoroughly and sensitively examine whether the hard>easy and easy>fix contrasts show similar sensory-biased organization, we correlated the vertices activations between both group average maps (i.e. **Figure 5b** and **Figure 6a**). Correlations between vertices for extended MD and Michalka regions combined showed a significant but weak negative correlation (*r* = -0.19; extended MD vertices only *r* = -0.25; Michalka-visual *r* = -0.48; but Michalka-auditory *r =* 0.3). These correlations, however, could arise due to the dependency between both contrasts. To de-correlate the contrasts, we repeated the analysis twice: once by splitting the subjects into two independent groups and once by correlating hard>easy with the average of the hard>fix and easy>fix maps. In both cases we again found a weak but significant negative correlation (*r* = -0.18 and -0.08 respectively for extended MD + Michalka regions). Even within an individual subject, correlating vertices of MD and Michalka regions between the two contrasts revealed no significant correlations (n=26) or weak negative correlations (n=10, range -0.03 to -0.2) and one subject with positive correlation (*r* = 0.1). These results suggest that there is little similarity in sensory-biased organization between the hard>easy and easy>fix contrasts.

Next, to quantify activation biases at the coarser region level, for each region we averaged the activation estimates (i.e. visual easy>fix minus auditory easy>fix) for all vertices and performed a one-sample t-test across subjects (p<0.05, Bonferroni corrected n= 64). The majority (41 out of bilateral 56) of extended MD regions showed significantly stronger visual than auditory activations (**Figure 7a)**. Among penumbra regions, the visual bias was strongest in dorsal parietal region LIPd, while among core regions, it was strongest in lateral frontal IFJp. 12 (out of the 36 bilateral) penumbra MD regions showed no overall visual vs auditory task preference (bilateral: s6-8, 6r, a32pr, SCEF; right: a10p, FOP5, TE1m, left: 11l) (**Figure 7a**). In the cases of 6r and SCEF, this is likely due to the antagonistic finer grained visual and auditory task biases within each region (**Figure 6a**). The only MD region with significantly stronger activation during the auditory task was left opercular penumbra region FOP5, just adjoining anterior insular region AVI, which itself had a stronger activation for the visual task.

**Figure 7.**
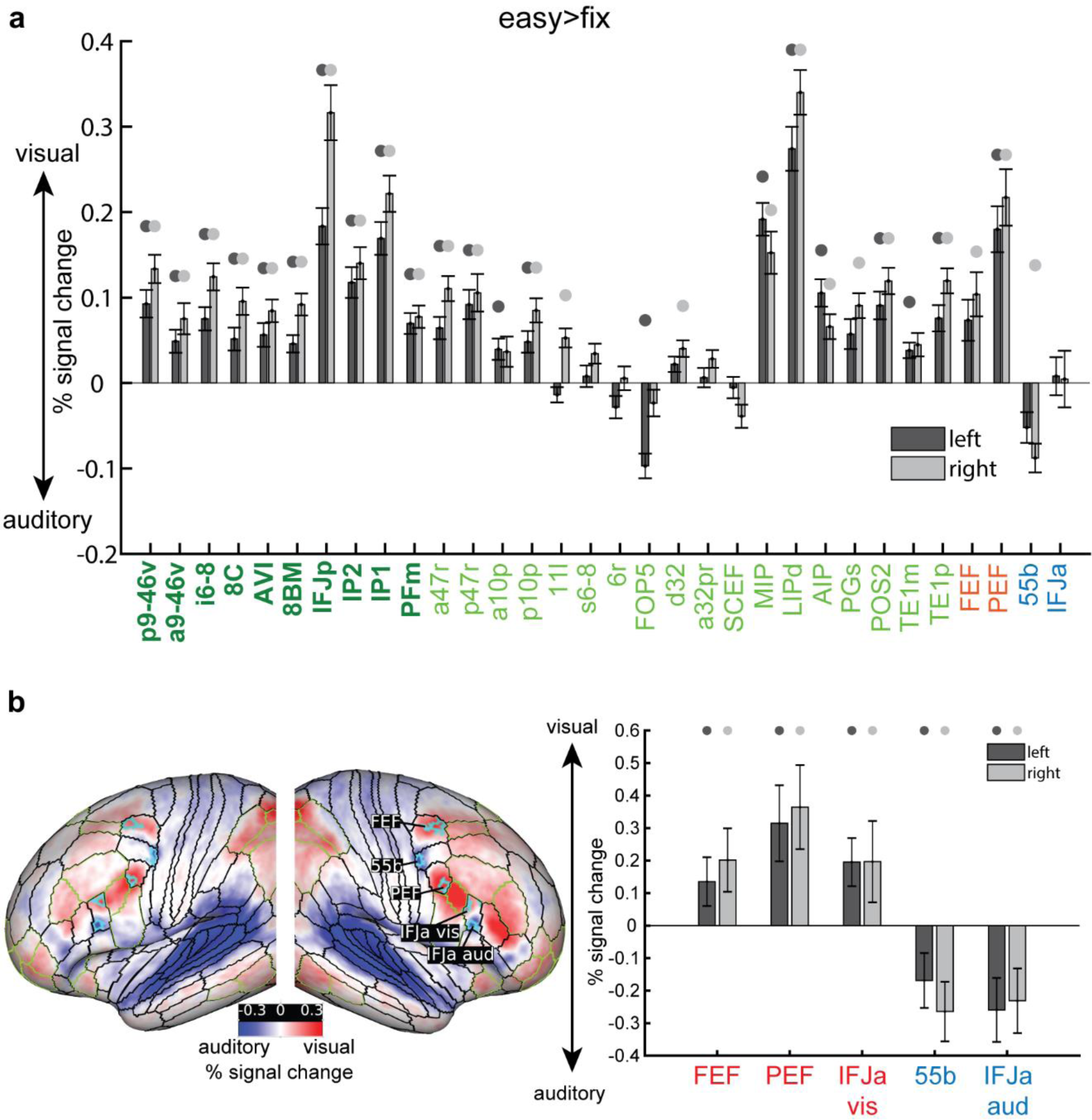
(a) Bar heights represent average activations (% signal change) for each region across subjects for the visual vs auditory easy>fix contrasts. Circles above bars denote p<0.05 Bonferroni corrected for n=64 regions. All remaining details are the same as in Figure 5a. **(b)** left: cyan borders surround the significant vertices (top 20%) within each of the HCP MMP1.0 regions overlapping with Michalka et al regions (Data is available at https://balsa.wustl.edu/NwV0p). Right: average activations (% signal change) for each region across subjects for the same contrast. Circles above bars denote p<0.05 Bonferroni corrected for n=10

Activations for MMP1.0 areas overlapping with the Michalka et al. (2015) regions also closely reflected the finer grained patterns in **Figure 6a**. Right FEF and bilateral PEF showed stronger visual than auditory task activations. Only right 55b showed stronger auditory than visual task activations (**Figure 7**). IFJa showed no overall task preference owing to its finer grained anatomical visual vs auditory split shown in **Figure 6a**. To address this issue, we repeated the above analysis for the Michalka et al regions using a finer-grained parcellation. To that end, we averaged two of the four runs of easy>fix visual vs auditory maps to select the top 20% (t-statistic) vertices showing visual bias within FEF, PEF and IFJa separately, and the top 20% (t-statistic) showing auditory bias within each of 55b and IFJa separately (**Figure 7b**). We then used the two remaining independent runs and averaged the easy>fix visual vs auditory activations for each group of vertices separately to get one value per region per subject. We then performed a one-sample t-test for each region against zero. Here, we found that all spatially constrained MMP areas overlapping with Michalka regions showed significant sensory biases as expected (**Figure 7b**). These findings are reliable as the results replicated with 10% and 30% vertices selected. These results highlight that the finest level of sensory-biased organization might not be well captured by the coarse level of MMP1.0 regions.

We also used these finer-grained regions to look again at the similarity of visual vs auditory task preferences in hard>easy and easy>fix contrasts. We again used independent runs to define the regions in the easy>fix contrast, and to estimate activations in the hard>easy contrast. Again, we found no statistically significant results for any of the smaller patches within the HCP MMP regions except for right 55b (p<0.05, Bonferroni corrected for 10 regions – 3 bilateral visual, 2 bilateral auditory). However, this was not a reliable finding and we observed null results for all regions when using the top 10% and 30% vertices.

It is also worth noting some findings in earlier cortical regions. Early auditory and visual regions showed stronger activations towards their respective tasks, which were more prominent and spatially extensive than revealed by the hard>easy contrast (**Figure 6a**). Within visual regions, foveal/central patches had stronger activations for the visual task while patches related to peripheral visual field showed stronger activations during the auditory task (**Figure 6a**). On the dorsal medial surface, we also identified several circumscribed regions with stronger activations during either visual or auditory tasks (**Figure 6a**).

These results paint a detailed anatomical picture regarding the cortical organization of MD and sensory biased regions. Instead of large smooth swathes of sensory biases along the lateral frontal cortex as identified by intrinsic rfMRI (see **Supplementary figure 3** for a direct comparison with rfMRI biases from Tobyne et al 2017), we found the lateral frontal surface decorated with multiple localized and interdigitated visual vs auditory task biases. For most MD regions, the easy>fix contrast showed a visual task bias, contrasting with highly similar visual and auditory responses for the hard>easy contrast. In line with the findings of Tobyne et al. (2017), several strong modality biases occurred just outside some core MD regions, including 8C, p9-46v and IP2.

### Sub-cortical and cerebellar MD sensory preferences

In this section, we investigated subcortical and cerebellar MD responses during the visual and auditory WM tasks. In our previous study (Assem et al. 2020), MD regions were identified in bilateral regions in the head of the caudate and in localized cerebellar regions (mainly cruses I and II) (**Figure 8a**). Their definition was based on a conjunction of co-activation during three cognitive demands and strong functional connectivity with the cortical core MD (Assem et al. 2020). Further putative bilateral anterior thalamic MD regions were also identified, though in this case based only on their strong functional connectivity with cortical core MD (**Figure 8a**). Here (see **Methods**) we used the rfMRI regions from Assem et al. (2020) to include the putative thalamic regions, but results for cerebellum and caudate were closely similar using the more spatially conservative task-based masks.

**Figure 8.**
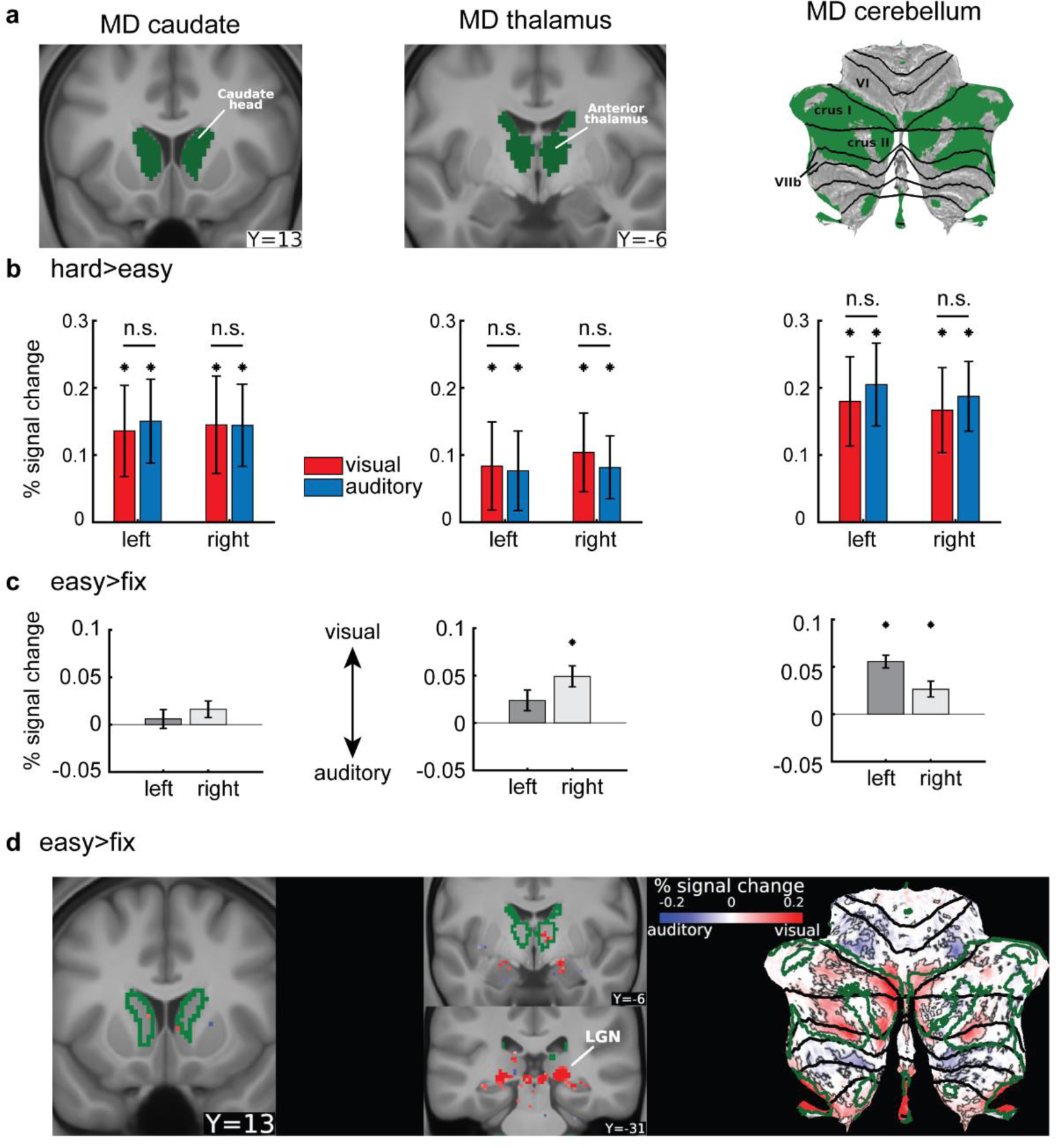
Subcortical and cerebellar MD visual vs auditory task preferences. **(a)** Subcortical and cerebellar MD masks identified in Assem et al 2020. The cerebellum is displayed as a flat surface with black contours representing anatomical borders. Data is available at https://balsa.wustl.edu/MxzxD **(b)** Hard>easy activations (% signal change) for the visual and auditory tasks. Left (MD caudate), middle (MD thalamus), right (MD cerebellum). Error bars are SEM. Asterisks denote significant hard>easy activations (p<0.05; Bonferroni corrected n=6 regions). n.s. denotes non-significant differences between visual hard>easy and auditory hard>easy (p<0.05; Bonferroni corrected n=6 regions). **(c)** Visual vs auditory easy task activations (% signal change) of subcortical and cerebellar MD regions (i.e. easy>fix visual – easy>fix auditory). **(d)** *Left and middle*: Subcortical voxels showing significant (FDR p<0.05) visual vs auditory task activations during easy>fix contrast. *Right*: Cerebellar voxels activations for the same contrast. Significant cerebellar regions are surrounded by white contours. MD regions are delineated by green contours. Data is available at https://balsa.wustl.edu/B4K41

First, we sought to confirm that the previously identified MD regions were activated during each of the visual and auditory hard>easy contrasts. For each region, we obtained a single estimate for the hard>easy activations (by averaging across all voxels within an MD region) and performed a one-sample t-test across subjects. Indeed, all bilateral caudate, thalamic and cerebellar MD regions showed significant hard>easy activations during both tasks (p<0.05, Bonferroni corrected for 6 regions; **Figure 8b**).

Next to unveil regions with statistically stronger activations for either the visual or auditory hard>easy contrast, we repeated the analysis previously used for cerebral cortex (i.e., for each subject we subtracted the auditory from the visual activations). First, we focused on the subcortical and cerebellar MD regions. For each MD region we obtained a single visual vs auditory value per subject then we performed a one-sample t-test across subjects. None of the MD regions showed stronger activation for either visual or auditory hard>easy contrasts (p>0.05, Bonferroni corrected for 6 structures; **Figure 8b**), similar to the cortical MD results.

Next, we repeated the same analysis using the easy>fix contrast (i.e. visual easy>fix minus auditory easy>fix). This contrast revealed stronger visual task activations in MD cerebellar regions bilaterally and the right MD thalamic region (p<0.05, Bonferroni corrected for 6 structures; **Figure 8c**). Interestingly, left MD thalamic and bilateral MD caudate regions failed to show any visual vs auditory task biases. These results again broadly align with the predominantly stronger visual activations in cortical MD during easy WM demands.

To also explore activations outside of MD regions and any finer grained patterns within MD regions, we performed a one-sample t-test on each subcortical and cerebellar voxel (p<0.05, FDR corrected). For the visual hard>easy vs auditory hard>easy contrast, we failed to identify any interpretable set of voxels either subcortically or in the cerebellum (not shown in a figure but available in the study’s BALSA files). However, for the visual easy>fix vs auditory easy>fix contrast, we identified a cluster of voxels in the posterior thalamus overlapping with the lateral geniculate nucleus (LGN) with visual preferences (**Figure 8b**). We also identified another set of voxels with auditory preferences immediately medial to LGN around the expected location of the medial geniculate nucleus (not shown in figure) which is a relay station for the auditory pathway. In the cerebellum, as expected, voxels within MD borders in cruses I and II (medial and lateral hotspots) showed stronger activations during the visual task. Medially, outside MD borders the stronger visual activations extended both dorsally (into lobule VI) and ventrally (into lobule VIIb), in line with previous studies identifying visual retinotopic responses in these regions (Brissenden et al. 2018; van Es et al. 2019). Laterally, MD borders were surrounded by patches of significant voxels with stronger auditory task activations both dorsally (lobule VI) and ventrally (lobule VIIb) (**Figure 8b**).

These subcortical and cerebellar results broadly mirror the cortical MD results. During easy cognitive demands (easy>fix), MD regions in the right thalamus and cerebellum showed relative preference for the visual task, while as cognitive demand increased (i.e. hard>easy), no significant task preferences were identified.

## Discussion

This study used the HCP’s multi-modal brain imaging methods to gain crisper resolution of the anatomical and functional organisation of domain-general MD and sensory-biased regions across the whole brain. Critically, we used two kinds of task contrast. Resembling many previous studies isolating MD activity, we used hard>easy contrasts in each modality. Resembling previous studies of modality-specificity, we used contrasts of a single task against a fixation baseline. Together, our findings give a comprehensive picture of the topographic organization of domain-generality and modality-specificity in the human brain.

### Near identical MD activations suggest similar visual and auditory integrative demands

For the hard>easy contrast, our results were clear-cut. Across the whole brain, this contrast produced almost identical activation for visual and auditory tasks. As anticipated, activation was focused on previously-identified MD regions; the brain-wide activation pattern, indeed, was strikingly similar to the MD pattern previously demonstrated, using different task contrasts, in HCP data (**Figure 3**). The strong similarity of visual and auditory results was demonstrable at the individual subject level (**Figure 3d**). It vividly illustrates the long sought-after anatomical precision in fMRI studies that is achievable by HCP-style projects. Additionally, such results point towards a way out of the reproducibility crisis in human brain imaging (Botvinik-Nezer et al. 2020) where a new approach to brain imaging acquisition, analysis, and data sharing (Glasser, Smith, et al. 2016) enables precise reproducibility of findings across studies. It is increasingly clear that, accompanying MD activity in cerebral cortex, a similar pattern of domain-general activity can also be seen in focal subcortical and cerebellar regions. In these regions too, our data showed no visual vs auditory task biases during the hard>easy contrast, further strengthening evidence for domain-general MD activity that extends across connected regions of cerebral cortex, caudate, thalamus and cerebellum.

MD regions are thought to play a core role in cognitive control (Cole and Schneider 2007; Power and Petersen 2013) through integrating different components to assemble a cognitive operation (Miller and Cohen 2001; Cole et al. 2013; Rigotti et al. 2013; Duncan et al. 2020). To solve the current tasks, for example, stimuli must be bound to their n-back positions, with more bindings to be reorganized for each new stimulus in the 3-back case. Our data confirm the domain-generality of MD activity, with identical response to similar control demands in visual and auditory tasks.

### Most MD regions showed stronger visual than auditory task activations

A different result emerged when easy tasks were compared to a no-task baseline. For most MD regions (cortical, subcortical and cerebellar), easy task versions showed stronger activations for the visual than for the auditory modality (easy>fix; **Figures 6**, **7**, **8**). This is despite the auditory task being harder, as indicated by behavioural accuracy. The visual task bias for MD regions broadly aligns with the Tobyne et al rfMRI findings (**Figure 1c**) and some previous task fMRI studies that did highlight visual preference for fronto-parietal activations (Braga et al. 2013; Mayer et al. 2016; Noyce et al. 2017). However, our anatomically refined results (**Figure 6)** extend these previous findings (see next section). The underlying mechanisms for this MD visual task bias remain unclear. Nevertheless, these results suggest MD regions are intrinsically biased to respond more strongly to visual than auditory tasks, perhaps reflecting the central role of visual processing in much primate - including human - cognition.

An exception for the stronger MD visual task activations is the left peri-insular area FOP5 (penumbra MD) which showed stronger activations during the auditory task, while the adjoining area AVI (core MD) showed stronger visual task activations. This observation fits with the broader literature on left peri-insular involvement in auditory, language and speech processing (Bamiou et al. 2003; Remedios et al. 2009). It is also in line with a recent invasive electrophysiology study in humans, which separated left opercular from anterior insular activity during a reading task by showing stronger responses in opercular electrodes (Woolnough et al. 2019). The fact that stronger auditory task activation within FOP5 was present only in the left hemisphere matches findings from a recent fMRI study, reporting articulation-related responses within the left MD anterior insula but not in the right hemisphere (Basilakos et al. 2018). Our study extends these findings by delineating these preferences between a penumbra and a core MD region.

Relatedly, a recent task fMRI study failed to find any sensory modality preferences in MD insular and anterior cingulate regions using a similar visual vs auditory 2-back>fix contrast (Noyce et al. 2017). In contrast, our results clearly highlight stronger activations during the visual task for 8BM (anterior cingulate) and AVI (insular) regions. One explanation for these conflicting results is our uncovering of adjoining regions with auditory task preferences near insula (AVI: visual, FOP5: auditory). Thus, a spatially coarse anatomical mask could mix signals across these functionally distinct regions and hide their task preferences. Further, Noyce et al. used a WM>baseline contrast to locate the cingulate region of interest. We have previously shown that task>baseline contrasts lead to posterior shifts in peak activations away from core MD regions (Assem et al. 2020). Thus, Noyce et al’s cingulate region likely focused on SCEF, just posterior to 8BM, which our current study showed had mixed finer grained modality preferences but no modality preferences at a coarse regional level.

In the cerebellum, previous studies identified both visual and auditory responses within cruses I and II, though without a clear delineation from MD regions (Petacchi et al. 2005; Kirschen et al. 2010; Brissenden et al. 2018; Ren et al. 2021). Here we show that MD portions of cruses I and II mirrored cortical MD findings by showing a dominant visual task preference. Subcortically, intriguingly, MD caudate regions did not show any task modality preferences during the easy>fix contrast. Only the right MD thalamic region showed stronger visual vs auditory task activations, copying the stronger right cortical MD visual task preference. It is worth noting that this right hemispheric dominance could be a feature of the n-back task. Language studies, for example, have shown that MD visual and auditory responses are stronger in the left hemisphere (Diachek et al. 2020).

For MD regions, in summary, there is a joint picture of predominantly visual bias when an easy task is compared to a no-task baseline, but precisely matched activity when contrasting hard>easy. Together, these results help synthesize prior indications of both modality preference and domain-generality

### Sensory biases surrounding MD cortex revealed during easy cognitive demands

The current study clearly separates MD regions from nearby regions with broadly stronger and more mixed sensory biases (**Figure 6**). On the lateral frontal surface, in addition to the visually biased MD region IFJp, we confirmed previous evidence that interdigitated auditory and visual patches lie immediately outside MD regions partially overlapping with regions FEF, 55b, PEF, IFJa (Michalka et al. 2015; Tobyne et al. 2017). In 55b, the auditory task biased patch was mostly localized to its posterior portion. However, it is worth noting that the topological organization of 55b is highly variable across individuals (Glasser, Coalson, et al. 2016) and the largely posterior activation likely reflects the consistency of the location of this portion across individuals. Additionally, there is topographic organization revealed with resting state functional connectivity along the posterior to anterior axis within area 55b (Glasser, Smith, et al. 2016; Van Essen and Glasser 2018) (Glasser et al., 2016; Van Essen and Glasser 2018). Our study’s high intrinsic spatial localization also enables separation of area IFJa into a ventral portion biased to respond more strongly during the auditory task and a dorsal portion biased towards the visual task. We found that these interdigitations also extend much more anteriorly than previously reported. Specifically, we identified three more interdigitated patches that partially overlap with IFSp (visual), IFSa (auditory) and penumbra MD region p47r (visual). We also found that the ventral (auditory)/dorsal (visual) division extends further anteriorly, where the peak of the visual task activation in IFSp is more dorsal while the peak auditory task activation in IFSa more ventral (**Figure 6a**). These partial overlaps between sensory-biased patches and HCP MMP1.0 areas demonstrate within-area heterogeneity in sensory-biased organization; however, it is as yet unknown whether such heterogeneity represents evidence for further areal subdivision, or if these multi-modal areas have topographically organized representation of sensory inputs (e.g., just as visual cortical areas separate upper and lower hemifields into different spatial locations, a multi-modal area might separate visual and auditory modalities into different spatial locations).

An interesting finding is that the spatial arrangement of these interdigitated regions, some of which harbour the strongest sensory biases, form an arc surrounding core frontal MD regions. These interdigitated regions are characterized by strong activations during multiple low cognitive demand contrasts (Assem et al. 2020). In some cases (e.g. FEF/PEF in current study; see **Figure 4**), further increase in demand leads to further increased activation, but this is by no means a general rule (Assem et al. 2020). While these demand-related activations could reflect activation bleeding from nearby MD regions (see **Figure 3**), they might also point to topographic extensions of MD properties. These “arc” regions also overlap with different resting-state networks (Ji et al. 2019; Assem et al. 2020). We have previously proposed that core MD regions lie at the heart of integrating different types of information fed in through surrounding regions (Duncan et al. 2020). In line with this proposal, the current anatomical arrangement suggests that interdigitated modality-biased regions might be important communication points between their affiliated resting-state networks and MD core.

In previous work, the neighbouring arrangement of MD and domain-specific regions (e.g. language) has been most clearly visible at the individual subject level (Fedorenko et al. 2012; Nieto-Castañón and Fedorenko 2012; Fedorenko and Blank 2020). In line with this, previous studies have shown that the frontal interdigitated sensory biases disappear in group-level maps (Noyce et al. 2017). Traditional group-level approaches heavily rely on cortical folds for inter-subject brain alignment, an approach limited by high inter-individual folding variability, especially in association cortices (Robinson et al. 2014, 2018; Coalson et al. 2018). Here, our method of inter-subject alignment (‘areal-feature-based’ MSMAll) uses multimodal structural and functional features that are more closely aligned with cortical areas, which significantly improves areal alignment (Robinson et al. 2014, 2018; Coalson et al. 2018). The result is that fine grained functional organization is visible at the group level (**Figure 6a**), allowing a straightforward and accurate comparison to findings from previous studies and to a canonical, high-resolution cortical parcellation (**Figure 6b**) (Assem et al. 2020).

Two more findings are worth noting. First, on the medial frontal surface a caudal-rostral division was visible in SCEF: its posterior portion showed stronger activations during the auditory task while its anterior portion showing stronger activations during the visual task. This result further supports the functional dissociation previously observed across SCEF (Assem et al. 2020) and broadly aligns with previous task fMRI indications of a spatially coarse caudal (auditory) to rostral (visual) medial frontal gradient (Mayer et al. 2016). Second, in early visual regions, peripheral visual regions showed a strong preference for the auditory task, while the visual task more strongly activated foveal/central regions. One possibility is that, as visual stimuli were foveal, the visual task enhanced activation of foveal regions, but suppressed peripheral regions. The data also align, however, with previous fMRI reports showing strong engagement of peripheral visual regions during auditory tasks (Cate et al. 2009) and aligns with anatomical evidence of direct connections between the primary auditory region and peripheral V1 regions (Falchier et al. 2002; Cappe and Barone 2005). Such results could indicate a functional link between peripheral visual regions and auditory processing, perhaps because auditory stimuli serve often to reorient gaze away from a currently foveated stimulus.

Finally, it is important to note two limitations in our study. First, because the visual and auditory tasks were performed on separate days and not within the same session, this might have weakened our statistical power to detect significant sensory biases in MD or Michalka et al regions for the hard>easy contrast. That said, the strong correlations between the visual and auditory hard>easy maps (**Figure 3**) make such an explanation unlikely. Even if any sensory biases existed, they are likely to be minute in comparison to the strong domain-general activations of MD regions. Second, because the visual and auditory stimuli in our tasks differed in multiple features, the relative sensory biases identified using the easy>fix contrast could reflect stimulus specific processing instead of modality preferences. For example, it has been previously argued that frontal visual-biased regions are more sensitive to spatial demands while auditory-biased regions are sensitive to temporal demands (Michalka et al. 2015). For this reason, it is important that the novel sensory biases we identified should be replicated using additional visual and auditory stimuli. Meanwhile, our replication of sensory-biased regions, identified in previous studies based on different tasks and task-free rfMRI (Tobyne et al. 2017), attest to the fitness of this contrast.

## Conclusion

Together, our results support the proposal of an integrative MD system, with some visual bias but a domain-general response to increased cognitive complexity. Adjacent to MD regions are interdigitated areas with visual and auditory preferences. Such regions are well-placed to feed modality-specific information into and out of the domain-general MD system. This arrangement may exemplify a more general motif, whereby domain-specific regions are placed to interact with domain-general processes of cognitive integration. The use of the HCP’s multimodal MRI acquisition and analysis approaches allowed this precision and replicability of results, paving a way out of the reproducibility crisis in neuroimaging, and opening the door to precise reference of imaging findings to a canonical, high-resolution cortical parcellation.

## Acknowledgments

J.D was funded by a Medical Research Council grant (SUAG/045.G101400); M.A was funded by Cambridge Commonwealth European and International Trust (Yousef Jameel scholarship); S.S. was funded by Gates Cambridge Trust (Cambridge, UK); M.F.G was funded by R01MH060974.

## Competing interests

The authors declare no competing interests

## Data availability

Data used for generating each of the imaging-based figures are available on the BALSA database (https://balsa.wustl.edu/study/x2m2L). Selecting a URL at the end of each figure will link to a BALSA page that allows downloading of a scene file plus associated data files; opening the scene file in Connectome Workbench will recapitulate the exact configuration of data and annotations as displayed in the figure.

## Supplementary figures

**Supplementary figure 1.**
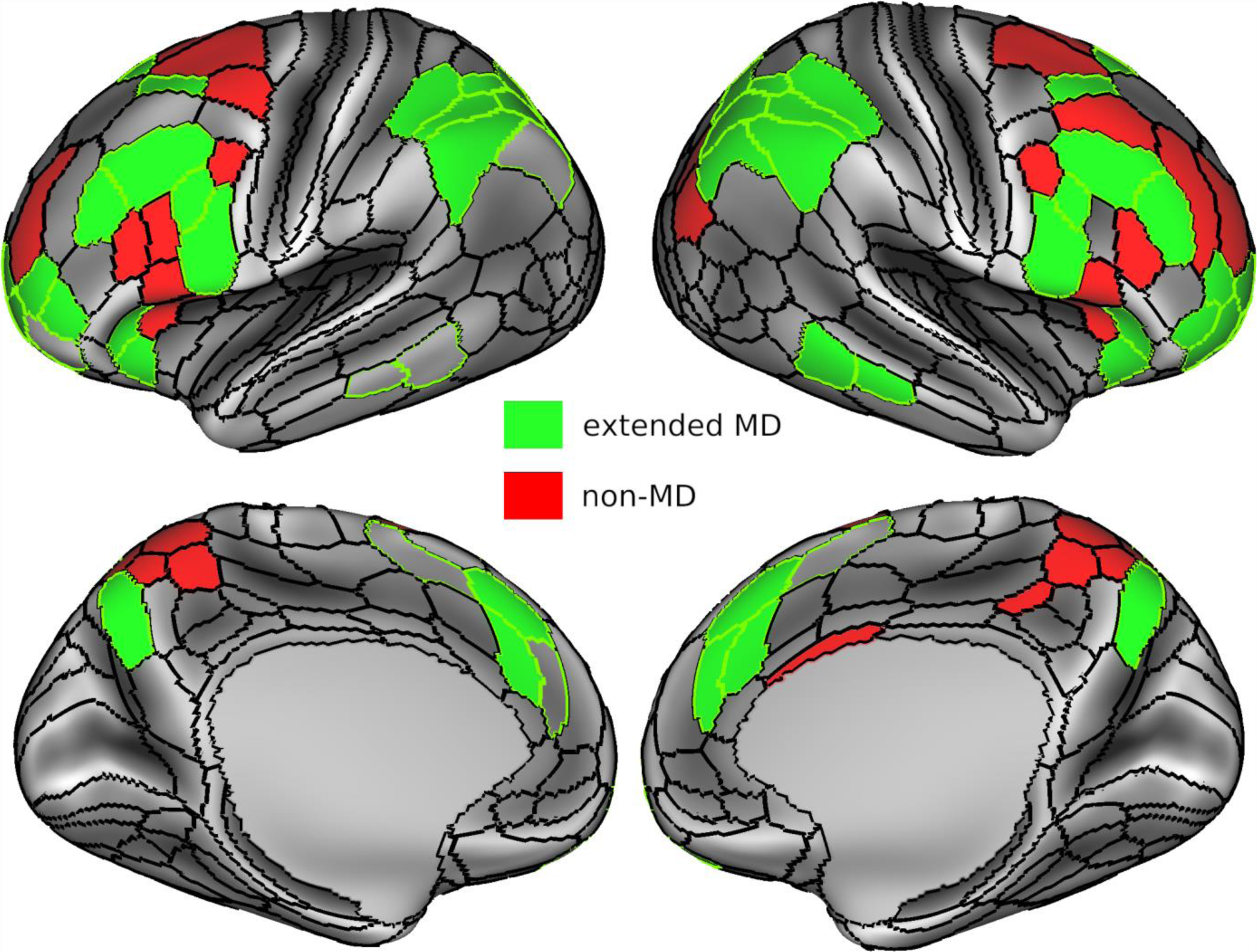
Conjunction of significantly activated parcels (p<0.05, Bonferroni corrected for 360 parcels) in both visual and auditory hard>easy contrast. Extended MD regions are in green, non MD regions are in red. Black contours belong to the borders of the HCP MMP1.0, green contours belong to borders of extended MD regions. Data is available at https://balsa.wustl.edu/G3l24

**Supplementary figure 2.**
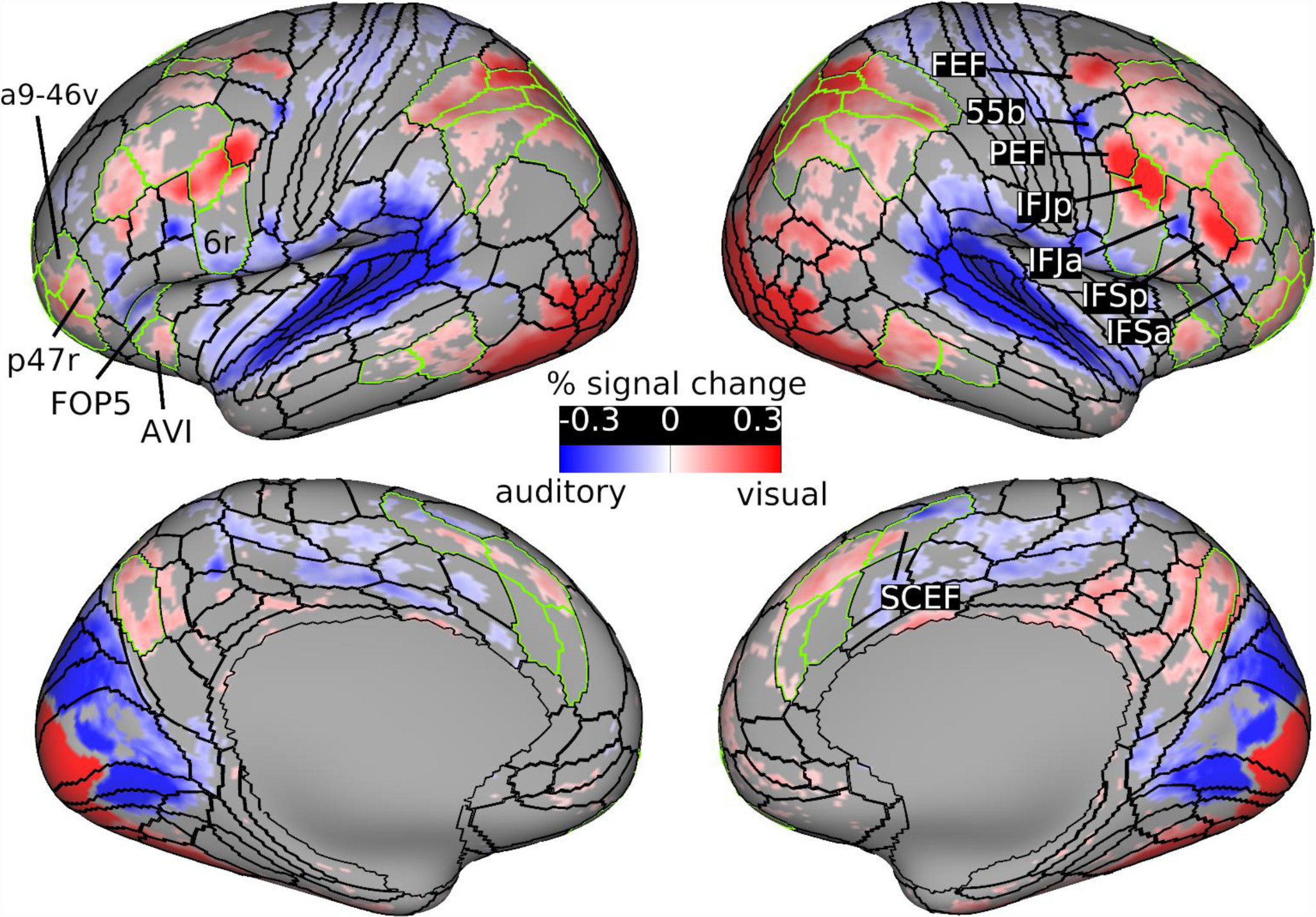
Significant vertices (colored) for the visual easy>fix minus auditory easy>fix contrast (FDR corrected p<0.05). Activations are % signal change. Black contours correspond to the HCP MMP 1.0 areal borders and green contours correspond to extended MD areal borders. Data is available at https://balsa.wustl.edu/zpj8K

**Supplementary figure 3.**
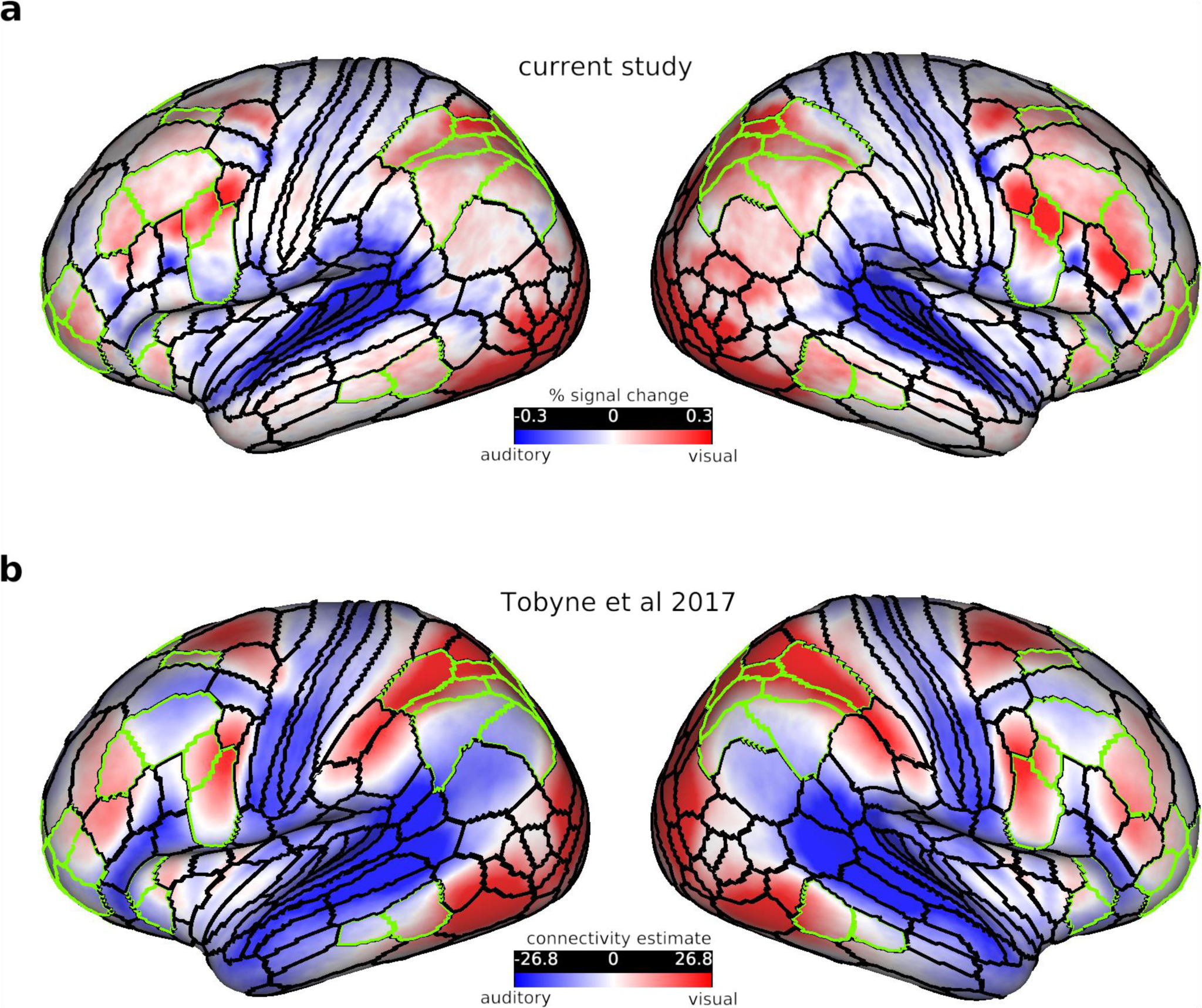
(a) easy>fix visual vs easy>fix auditory activations from current study. (b) Auditory vs visual modality biases as defined by Tobyne et al (2017) using rfMRI connectivity (see Tobyne et al 2017 for details on methods). Black contours correspond to the HCP MMP 1.0 areal borders and green contours correspond to extended MD areal borders. Data is available at https://balsa.wustl.edu/X5Pjj

## Notes

https://balsa.wustl.edu/study/x2m2L

### Competing Interest Statement

The authors have declared no competing interest.

### Summary of Updates

More analysis to probe any minute sensory-biased organisation in the hard>easy contrast. None were found. Also figures colour scales and labels are updated to make them clearer.

